# Decorrelation Time Mapping as an Analysis Tool for Nanobubble-Based Contrast Enhanced Ultrasound Imaging

**DOI:** 10.1101/2022.12.21.521428

**Authors:** Dana Wegierak, Michaela Cooley, Reshani Perera, William J. Wulftange, Umut A. Gurkan, Michael C. Kolios, Agata A. Exner

## Abstract

Nanobubbles (NBs) are nanoscale (∼100-500 nm diameter) ultrasound (US) contrast agents that enable new robust applications of contrast enhanced US and US-mediated therapy. Due to their sub-micron size, high particle density, and highly deformable shell, NBs exhibit unique properties. In pathological states of heightened vascular permeability, such as in tumours, NBs can extravasate, enabling extravascular applications not currently possible with clinically available microbubbles (∼1000-10,000 nm diameter). This ability can be explored to develop imaging biomarkers to improve tumour detection. There is a need for an imaging method that can rapidly and effectively separate intravascular versus extravascular NB signal when imaged using nonlinear dynamic contrast enhanced US. Herein, we demonstrated the use of decorrelation time (DT) mapping to achieve this goal. Two *in vitro* models were used to explore the roles of NB velocity and diffusion on DTs. Mice bearing prostate specific membrane antigen (PSMA) expressing flank tumours (n = 7) were injected with bubble agents to evaluate the *in vivo* potential of this technique. The DT was calculated at each pixel of nonlinear contrast videos to produce DT maps. Across all models, long DT correlated with slowly moving or entrapped NBs while short DT correlated with flowing NBs. DT maps were sensitive to NBs in tumour tissue with high average DT in tumour regions (∼10 s) compared to surrounding normal tissue (∼1 s). Molecular NB targeting to PSMA extended DT (17 s) compared to non-targeted NBs (12 s), demonstrating sensitivity to NB adherence dynamics. Overall, DT mapping of *in vivo* NB dynamics produced detailed information of tumour tissue and showed potential for quantifying extravascular NB kinetics. This new NB-contrast enhanced US-based biomarker can be useful in molecular ultrasound imaging, with improved sensitivity and specificity of target tissue detection and potential for use as a predictor of vascular permeability and the enhanced permeability and retention (EPR) effect in tumours.

## Introduction

Tumours are characterized by high vascular permeability and low lymphatic drainage, which increases the potential for enhanced permeability and retention (EPR) of macromolecules (∼200 nm).^1,2^ Macromolecular substances with high biocompatibility circulate in the blood pool for extended periods and remain there in the presence of healthy tissue and blood vessels.^1^ During their circulation, however, these macromolecules selectively leak out of tumour blood vessels due to disorganized and pathological vascular wall structure.^1^ Low lymphatic drainage limits the exit of macromolecular substances from the tumour parenchyma, leading to their accumulation. When present, EPR leads to the preferential tumour-localized accumulation of nano-agents by up to a factor of two over background.^3^ As such, understanding and characterizing the EPR effect is a key motivation for nanomedicine-based agents in oncology for both diagnostic and therapeutic applications.^4^

Cancer diagnosis relies heavily on biomedical imaging technologies, like magnetic resonance imaging, positron emission spectroscopy, or ultrasound (US).^5^ To increase specificity and sensitivity of detection, molecular imaging can be utilized, which employs agents specifically designed to target biomarkers overexpressed on cancer cells (e.g., antibody conjugation to nano-agents).^3,4,6^ Molecular imaging using PET/SPECT, MRI, and optical imaging has played a significant role in cancer diagnosis with many new agents either in clinical trials or in robust preclinical development.^7–11^ Although all of these techniques represent important advances toward efficient and sensitive tumour diagnosis, they also contribute to the high cost (expensive imaging technology) and risk (ionizing radiation) of nanotherapeutics in oncology. These barriers are exasperated by the fact that in the clinical setting, target tumour epitopes may be unidentified during the early stages of diagnosis, requiring tissue biopsy, which is limited when tumours are small. Previous groups have shown that when a human tumour expresses the EPR effect, nanotherapeutics have had notable success.^12,13^ Quantitation of EPR has therefore been identified as a critical element for the success of targeted molecular techniques for cancer diagnosis and treatment.^6,13^

US is safe and low-cost relative to other imaging technologies.^14,15^ The use of contrast agents in combination with nonlinear contrast (NLC) imaging produces detailed images and is FDA approved in clinical practice for echography and imaging of liver lesions.^16^ US contrast agents are typically lipid-stabilized bubbles with perfluorocarbon gas cores ranging from 1 to 10 µm in diameter (microbubbles - MBs).^16,17^ Based on the acoustic properties of gas-filled MBs, binding specific probes to the MBs allows targeting to histological structures of interest and is termed US molecular imaging.^18^ US molecular imaging is an ongoing area of interest that has garnered significant attention in cancer detection.^19–22^ However, due to the size and mode of delivery, MBs for US molecular imaging mainly target intravascular epitopes.^23^ Thus, they offer minimal information about tumour microenvironment characteristics and cannot be used to target biomarkers overexpressed on the cancer cells themselves.

Nanobubble (NB) US contrast agents (∼100–500 nm diameter), unlike their larger counterparts, are not restricted to the blood pool when in the presence of pathologically leaky vasculature.^24–27^ NB formulations previously developed by our group^28^ are compatible with existing nonlinear US contrast imaging modes and feature high stability (∼18 min half-life), while being easy to adapt for molecular targeting.^27^ These NBs exhibit prolonged retention in tumours and exhibit a large area under the curve (AUC) in contrast enhanced US time–intensity curves.^29^ However, currently available post processing methods cannot effectively distinguish signal of extravascular nanobubbles from blood pool populations.

Decorrelation of US echo between subsequent frames has previously been studied^30^ and is a method of analysis being explored for thermal^31^, radiofrequency^32,33^, and microwave^34^ ablation therapy monitoring and assessment. Echo decorrelation has also attracted attention for blood velocimetry.^35,36^ Blood cells travelling through an intravascular ultrasound imaging plane generate echo signals which decorrelate at a rate related to the scatterer flow velocity.^35,36^ It is generally known that the movement of blood, and therefore blood-pool agents, causes fast decorrelation of the ultrasound signal. For instance, it has been shown that MBs decorrelate from frame to frame on the millisecond timescale.^37^ For these reasons, high frame rate US imaging (>500 fps) of MBs (blood-pool agents) using US echo decorrelation has been useful in US localization microscopy, where the signal from a blood pool agent is tracked to recover the vascular structure from the flow pathway.^37,38^ High frame rates, however, contribute to not only rapid agent destruction, but also large datasets requiring robust memory solutions. Assessment of echo decorrelation at low frame rates would reduce bubble destruction, reduce memory storage requirements, and should therefore be explored further. Decorrelation of US echo signals has been used to understand the effects of blood and its components on the acoustic response of NBs.^39^ What remains is the need to adapt this 1-D signal analysis method to US videos as a a bridge to rapid signal assessment with spatially localized trends.

The following work aims to investigate the principles of echo decorrelation for slow moving and stationary agents for improved cancer detection by the identification of localized agent accumulation. Herein, we present a decorrelation time (DT) mapping technique to simultaneously assess the morphology and physiology of prostate cancer tumours through the US signal analysis of NB motion with broad applicability across vascular diseases that result in increased vascular permeability. While we do not resolve individual NBs, we track the change in NLC mode US intensity over time as a measure of NB motion within the resolution volume of the ultrasound imaging device at low frame rates.^40^ We validated the DT technique in two *in vitro* phantom experiments to show the ability of the DT mapping method to distinguish slow and rapid moving agents in the presence and the absence of permeation. DT mapping was then applied to contrast-enhanced US data of mouse flank tumour models filling with NBs compared to microbubbles. Overall, DT mapping is a robust imaging technique for molecular ultrasound image processing to detect and characterize NB-based biomarkers of contrast-enhanced ultrasound imaging in tumours.

## Methods

### Preparation of Nanobubbles

Lipid shell stabilized C_3_ F_8_ NBs were formulated as previously described.^28,41^ Two forms of NBs were used: 1) plain NBs – for passive targeting and accumulation dynamics - and 2) prostate specific membrane antigen targeted NBs (PSMA-NBs) – for active targeting and enhanced retention at the tumour site.^41^ Agents were activated by mechanical agitation (Vialmix - Bristol-Meyers Squibb Medical Imaging, Inc.; N. Billerica, MA) and isolated by differential centrifugation. NBs and PSMA-NBs had average diameters of 284± 96 nm and 277± 97 nm respectively,^39^ as measured using resonance mass measurement instrumentation (Archimedes, Malvern Panalytical Inc., Westborough, MA).^41,42^

### Preparation of Microbubbles

For *in vitro* studies, microbubbles (MBs) with the same lipid shell composition as the NBs were prepared according to Abenojar et al.^43^ Briefly, the MBs were isolated by diluting the bubble solution obtained after mechanical agitation in 10% vol/vol glycerol/propylene glycol in phosphate buffered saline (PBS). Following dilution, the polydisperse bubbles were centrifuged at 300 g for 10 min and the infranatant was discarded. The remaining MB solution was re-dispersed in PBS and centrifuged at different sequential speeds (30, 70, 160, and 270 g) for 1 minute each, after which the infranatant was collected. The desired MB sample (mean diameter = 875 ± 280 nm, mode diameter = 1µm, concentration = 1.18 × 10^10^ bubbles/mL) was then transferred to a 3 mL headspace vial, capped with a rubber septum, and sealed with an aluminum cap.^43^ The vial was then flushed with 10 mL of C_3_ F_8_ gas and used immediately for imaging. For *in vivo* studies, commercially available MBs (Lumason®, Bracco Diagnostics Inc.) were used for comparison to NBs. Lumason is globally known as SonoVue® and has a mean bubble diameter of 2.5 µm with more than 90% of the bubbles smaller than 8 µm.^44^ Lumason® was prepared according to the protocol provided by the manufacturer.

### Image Acquisition Protocols

US imaging for all models was conducted using the Vevo 3100 (VisualSonics) US system using an MX250 transducer (15–30 MHz) with an 18 MHz center transmit frequency linear array transducer in dual B-mode and RF-nonlinear contrast mode (RF-NLC). NLC imaging on the Vevo 3100 has been implemented using pulse amplitude modulation. Amplitude modulation uses multiple pulse firings to differentiate linear and nonlinear US signals. For ease of comparison, the imaging parameters for each setup are summarized in Table 1.

**Table 1.** Model imaging conditions.

| <u>Model/Setting</u> | <u>Flow Phantom Model</u> | <u>Microfluidic Model</u> | <u>Mouse Model</u> |
| --- | --- | --- | --- |
| *Center Frequency* | 18 MHz | 18 MHz | 18 MHz |
| *Dynamic range* | 40 dB | 40 dB | 40 dB |
| *Gain* | 30 dB | 30 dB | 30 dB |
| *Beamwidth* | Medium | Medium | Medium |
| *Transmit Power* | 4% | 20% | 4% |
| *Frame rate* | 20 fps | 5 fps | 5 fps |
| *Number of frames* | 100 | 4000* | 1000 |

\*Analyzed 200 frames at a time for decorrelation time mapping

### Validation of Decorrelation Time in a Flow Phantom

To formulate an *in vitro* model featuring controlled regions of stationary agents and moving agents, a simple flow phantom was produced (schematic shown in Fig. 1.) through adaptation from a previously established protocol.^40^ Low temperature gelling agarose (Sigma Aldrich, USA) was prepared in 1.25% w/v concentration in PBS in a 100 mL beaker and heated in the microwave until the agarose powder was completely dissolved. The mass of the beaker with agarose and PBS was measured before and after heating. Distilled, deionized (DDI) water was used to compensate for loss by evaporation during the heating process. After compensatory addition of DDI water, the beaker solution was covered by a plastic film to minimize additional evaporation and set aside to cool to 40 ^°^ C (congealing temperature 26–30 °C). Upon reaching 40 ^°^ C, NBs (∼4.07×10^9^ NBs/mL) were mixed into the agarose gel solution and poured into a phantom holder and allowed to set around a needle (0.51 mm O.D.) at room temperature (22 °C). When gelling was complete, the needle was removed and the same concentration of NBs as in the gel were perfused through the channel at 250 µL/min during imaging. From previous studies, the average pore size from 1.25% w/v concentration agarose matrix is ∼340 nm.^40^ Flow phantoms were produced immediately prior to imaging.

**Figure 1.**
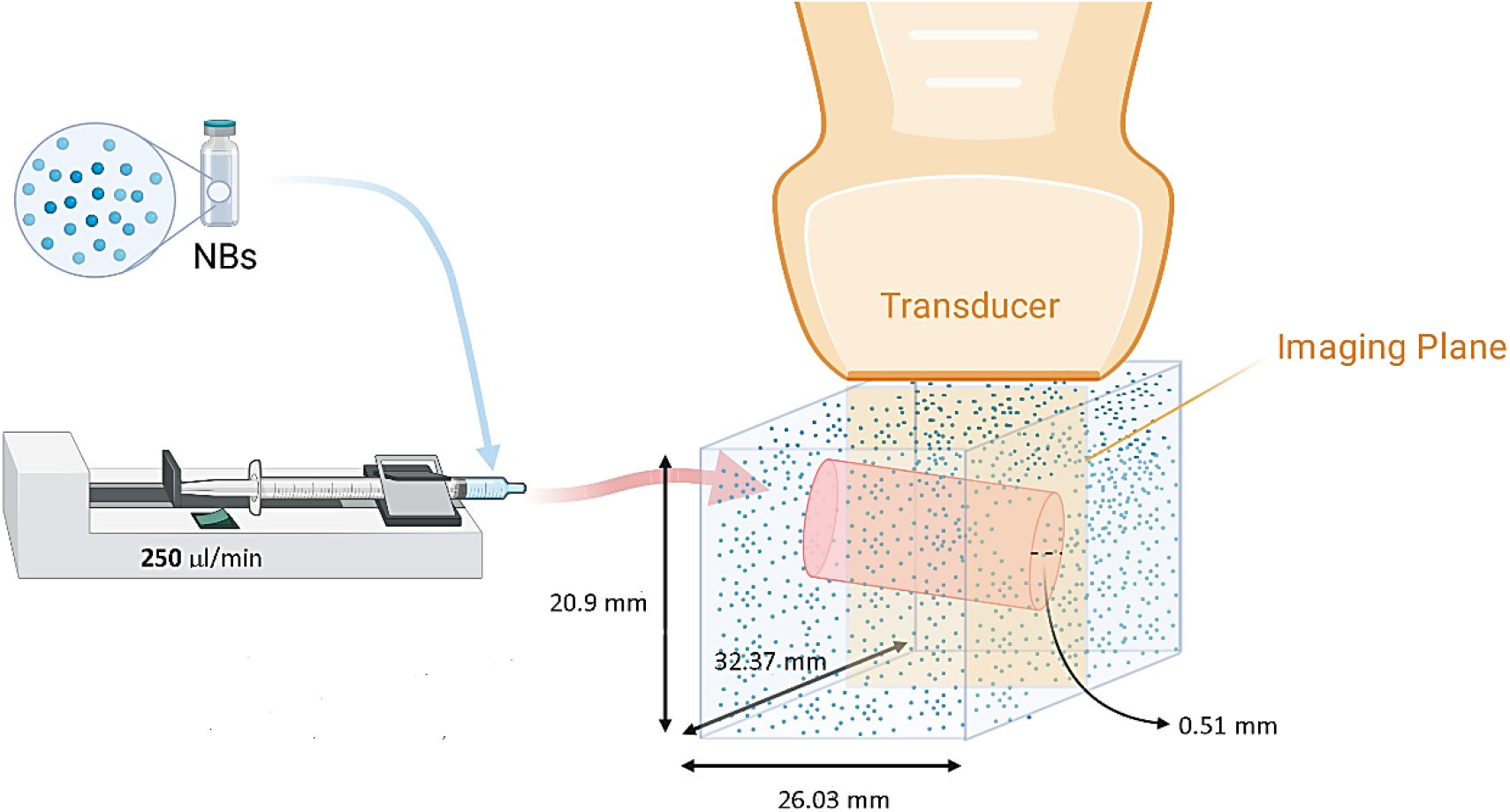
Schematic of a (1.25%) agarose gel phantom with NBs (100-500 nm diameter; concentration ∼4×10^9^ NBs) entrapped in the agarose matrix (∼340 nm pore size). During imaging, a dilution of NBs in PBS was produced to match the concentration of entrapped agents and flown through the hollow channel at a rate of 250 µL/min using a syringe pump F100X (Chemyx Inc. North Americ, Stafford, TX).

For the agarose flow phantom setup, the US transducer was coupled to the phantom using PBS. The focus of the transducer was aligned to the center of the flow channel. Phantoms were imaged at 20 frames per second (fps) with a contrast gain of 30 dB, B-mode gain of 28 dB, dynamic range of 40 dB, medium beamwidth and 4% transducer power (system minimum) for a total of 100 frames. All 100 frames were analyzed at a time for Decorrelation Time mapping.

### Validation of Decorrelation Time in a Microfluidic Flow Phantom

As an intermediate *in vitro* model with progressive extravasation and progressive entrapment of agents, a microfluidic chip with collagen I matrix (Advanced Biomatrix, Carlsbad, CA) surrounding a lumen was fabricated (schematic shown in Fig. 2.) as previously described.^45^ The device, LumenChip, contained an extracellular matrix (ECM) chamber with a lumen extending through the center. The lumen was formed by casting a collagen I solution around a needle and allowing the collagen to polymerize before removal of the needle. Collagen I matrices were produced to yield average pore sizes of 1.3 ± 0.7 μm or 3.7 ± 1.3 μm in diameter.^46^ Difference in average matrix pore size allowed for comparison of extravasation from the luminal space. To assemble the chip, double-sided adhesive (3M, Saint Paul, MN) and polymethyl methacrylate (McMaster-Carr, Elmhurst, IL) were layered to create a chamber with an inlet and outlet through which a needle (0.5 mm outer diameter; Component Supply Company, Mentor, OH) could be inserted. The top and bottom layers of the device were made of Sylgard™ 184 polydimethylsiloxane (PDMS) (The Dow Chemical Company; Midland, MI). The PDMS was cast (9:1 ratio) and cured overnight on a leveling plate to achieve a thickness of ∼1.5 mm PDMS.

**Figure 2.**
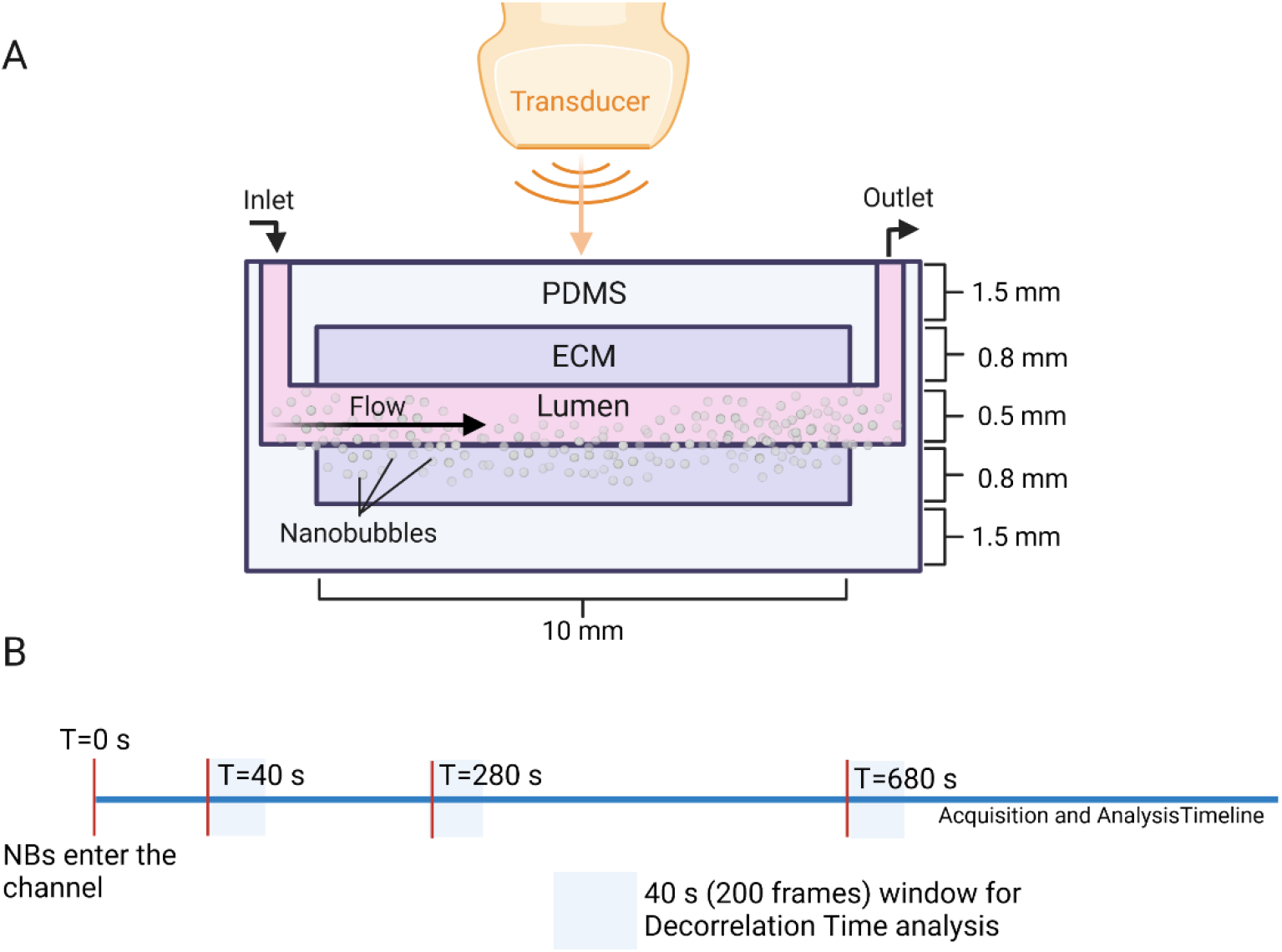
A) Schematic representation of microfluidic setup under ultrasound. The microfluidic device contains an extracellular collagen I matrix chamber with a lumen formed by polymerization around a needle (0.5 mm), which extended through the center of the device. The collagen I matrix (ECM) was produced with either 1.3 µm or 3.7 µm average diameter pore sizes.^46^ B) Data timeline showing 4000 frames of imaging and the 200-frame intervals that were selected for analysis by decorrelation time mapping.

During imaging, 1 mL PBS with either NBs (2.03 × 10^10^ NBs/mL) or MBs (1.18 × 10^7^ MBs/mL) was perfused through the channel at 50 µL/min using a Fusion 200-X syringe pump (Chemyx; Stafford, TX). The US transducer was coupled to the device using water. The focus of the transducer was aligned to the bottom wall of the microfluidic device, below the lower boundary of the matrix. LumenChips were imaged using the FUJIFILM VisualSonics Vevo 3100 at 5 fps with a contrast gain of 30 dB, B-mode gain of 28 dB, dynamic range of 40 dB, medium beamwidth and 20% transducer power. Devices were imaged for 4000 frames, starting just prior to perfusion, and capturing the agent dynamics from the start of perfusion through diffusion and extravasation. Intervals of 200 frames were selected from these 4000-frame videos starting at 40, 280 and 680 s after agent entry into the lumen for subsequent analysis by Decorrelation Time mapping.

### Application of Decorrelation Time in nanobubble dynamic Contrast Enhanced US in Tumour-Bearing Mice

The *in vivo* NLC data was obtained from a previously published study.^41^ A simplified reiteration of the protocol is briefly summarized here; for in-depth detail, the reader is directed to the original publication. Seven mice bearing subcutaneous human PC3-pip tumour xenografts were analyzed. MBs, NBs or PSMA-NBs were prepared as described above and in the published study and were injected via the tail vein in 6 mice (Fig. 3-A). PSMA is a membrane bound glycoprotein that is highly expressed on the cell surface of prostate cancer cells and has previously been used by our group as a targeting epitope for PC3pip tumours.^27^ For the seventh mouse, PSMA-NBs were injected via tail vein and imaged for the study of the dynamics of targeted NBs (PSMA-NBs). Next, a burst sequence was delivered to the imaging plane, followed by a 30 min period with no imaging, to eliminate any PSMA-NB agents from the tumour and blood circulation. While maintaining the transducer alignment on the mouse, NBs were then injected to the same mouse via the tail vein and imaged (see schematic Fig. 3-B).

**Figure 3.**
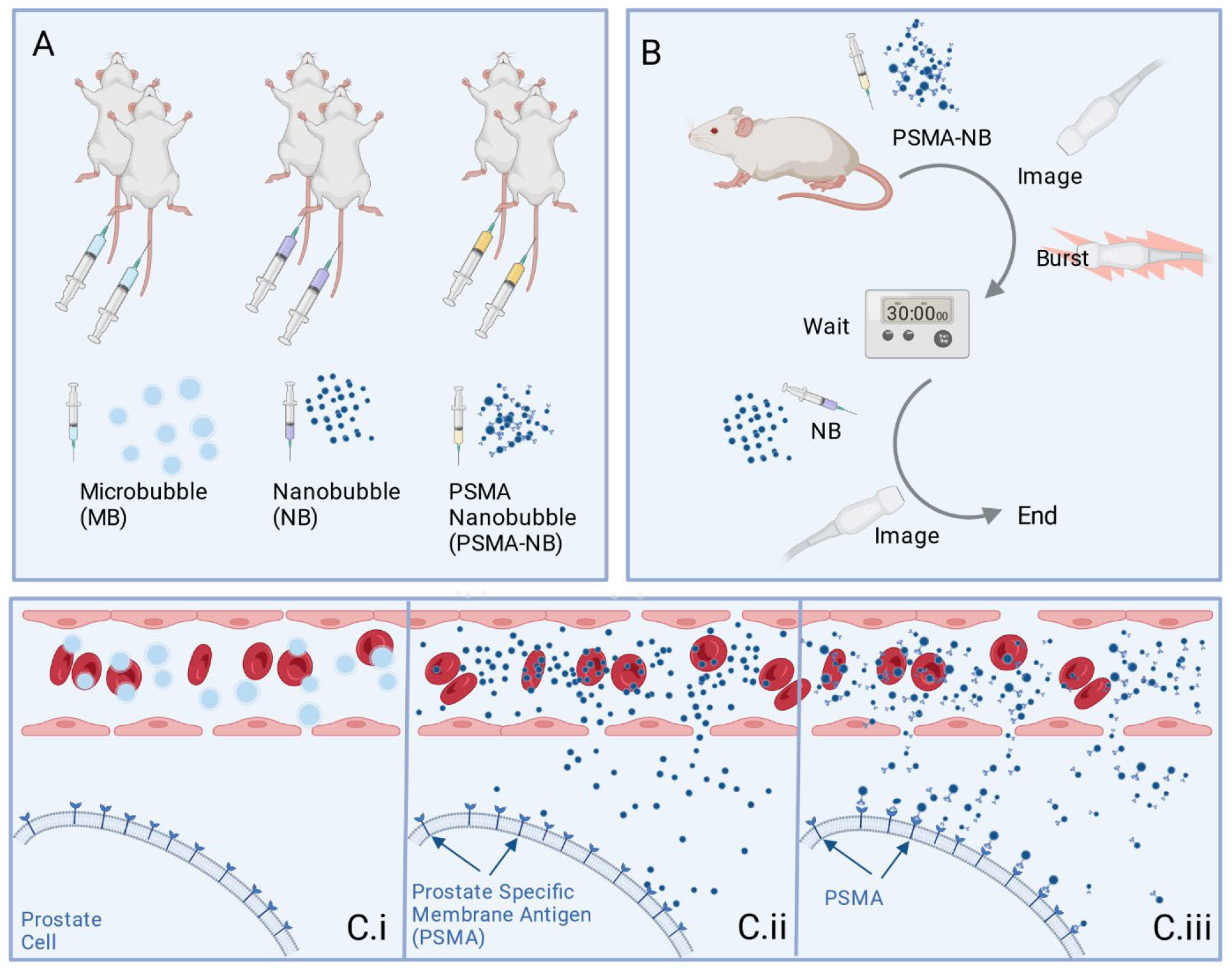
A) Depiction of flank tumour mouse models injected via tail vein with either microbubbles (MBs; Lumason), nanobubbles (NBs), or prostate specific membrane antigen targeted NBs (PSMA-NB). B) Workflow for a single flank mouse tumour model injected and imaged with PSMA-NBs followed by plain NBs. C.i) MBs are blood pool agents, restricted to the vasculature. C.ii) NBs can penetrate past endothelial gaps of leaky vasculature. C.iii) Molecular targeting is used to enhance retention of nanobubble agents at the surface of prostate cancer cells.

For the *in vivo* mouse setup, the US transducer was coupled to the subcutaneous tumour with US gel. The focus of the transducer was aligned to the bottom edge of the tumour. Tumours were imaged at 5 fps with a contrast gain of 30 dB, B-mode gain of 28 dB, dynamic range of 40 dB, medium beamwidth and 4% transducer power. Agent kinetics *in vivo* were measured with NLC mode starting from baseline and including agent injection for a total of 1000 frames. The first frame of the NLC *in vivo* video was used for baseline subtraction by using image registration (MATLAB - *imregister()*) to subtract residual tissue signal and mitigate motion artifacts. The final matrix was then analyzed via Decorrelation Time mapping.

### Decorrelation Time Mapping

The autocorrelation function is used in time series analysis to identify temporal trends. The time at which the autocorrelation function decorrelates to a set threshold is known as the decorrelation time (DT) and can be used as a measure to represent the rate of decorrelation and the strength of the trend in the time series. DT was defined as the time shift required for the correlation coefficient to reach 0.5, similar to Jeon at al.^39,47^ NLC mode US videos of each model were captured and stored as real-valued matrices (*m*×*n*×*t*) (*m* - image depth, *n* - image width, *t* - number of frames collected at frame rate, *fs*; matrix depiction in Fig. 4-A.) in MATLAB for analysis. A pixel time-intensity curve (TIC) was selected (*m*_i_,*n*_i_ at all *t* and normalized by z-normalization (MATLAB function, *normalize(, ’zscore’)*). The intensity autocorrelation function for each normalized pixel TIC was calculated using the MATLAB function, *xcorr()*^48^. The autocorrelation function was converted from a function of lag to a function of time using the frame rate (*fs*) of the video. The time at which the autocorrelation function of the pixel TIC decayed to 50% was used as the DT at that spatial location. This process was iterated over every spatial location to produce the final map, as depicted in Fig. 4.

**Figure 4.**
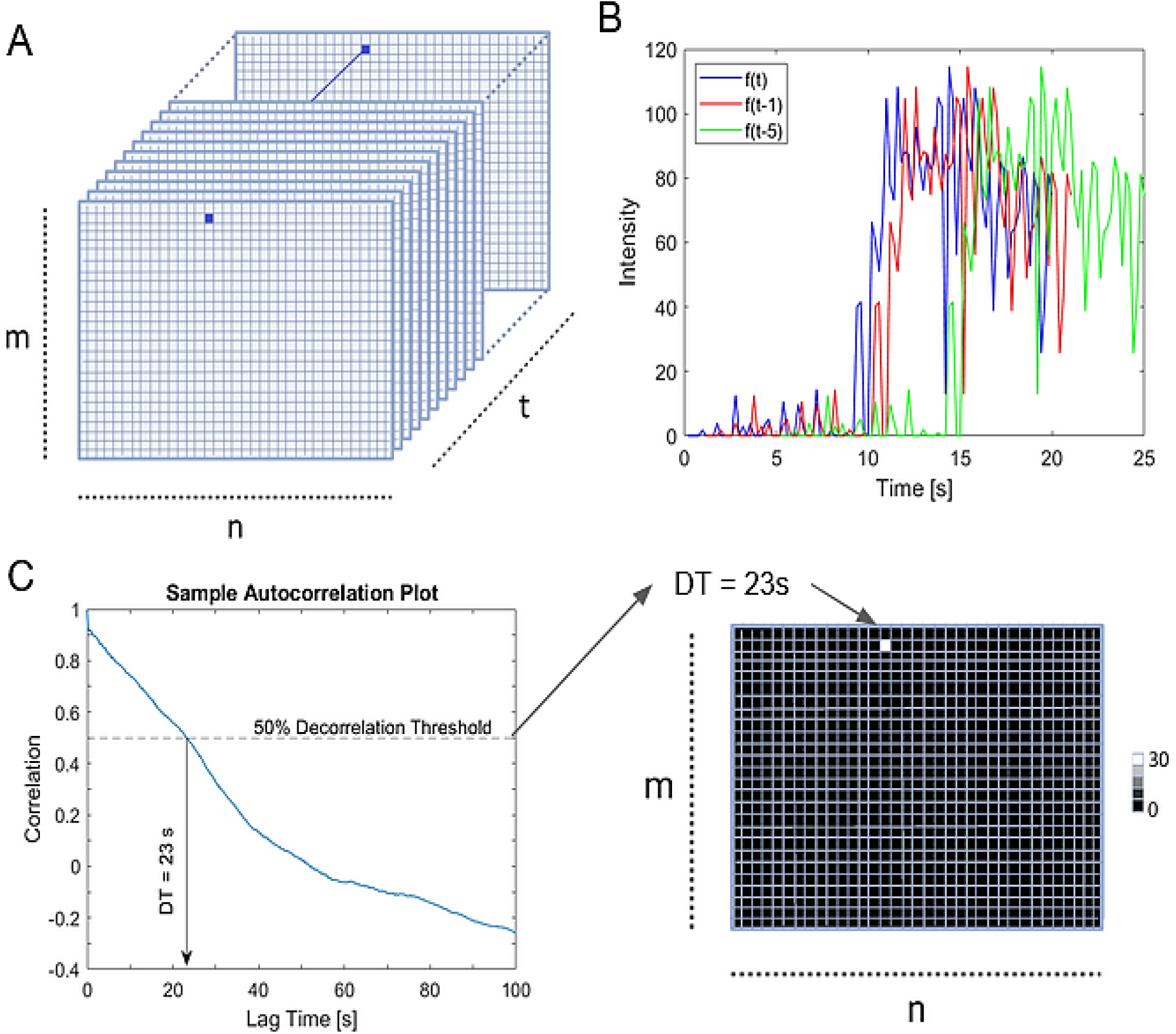
A) Depiction of a data matrix from imaging videos where the ROI is *m* x *n* pixels with *t* frames. The blue square indicates a selected pixel used to produce a time-intensity curve (TIC). B) Sample TIC f(t) from pixel *p(m,n)* for all frames with the representative TICs after time lag ‘x’ - (*f(t-x)*). C) Sample autocorrelation plot; the dashed line indicates the 50% decorrelation threshold, and the solid line arrow indicates the decorrelation time. The decorrelation time is mapped to a new matrix -*DT(m,n)*-that contains the decorrelation times corresponding to each pixel *p(m,n)*.

## Results

### Flow Phantom

A simple flow phantom was produced featuring well-defined regions of stationary and flowing agents that could be imaged in the same field of view using NLC mode US imaging. Representative images of signal intensity for the first and last NLC frames of the flow phantom experiment with entrapped NBs are presented in Fig. 5-A&B. The center channel with flowing agents was visibly difficult to distinguish from the surrounding, entrapped agents in the static NLC images (Fig. 5-A). The bottom surface of the channel was visible in the last frame of imaging due to localization of agents at the channel wall, likely caused by acoustic radiation force (Fig. 5-B). The signal in the surroundings also decreased over time because of the dissipation of entrapped NBs from extended US exposure.

**Figure 5.**
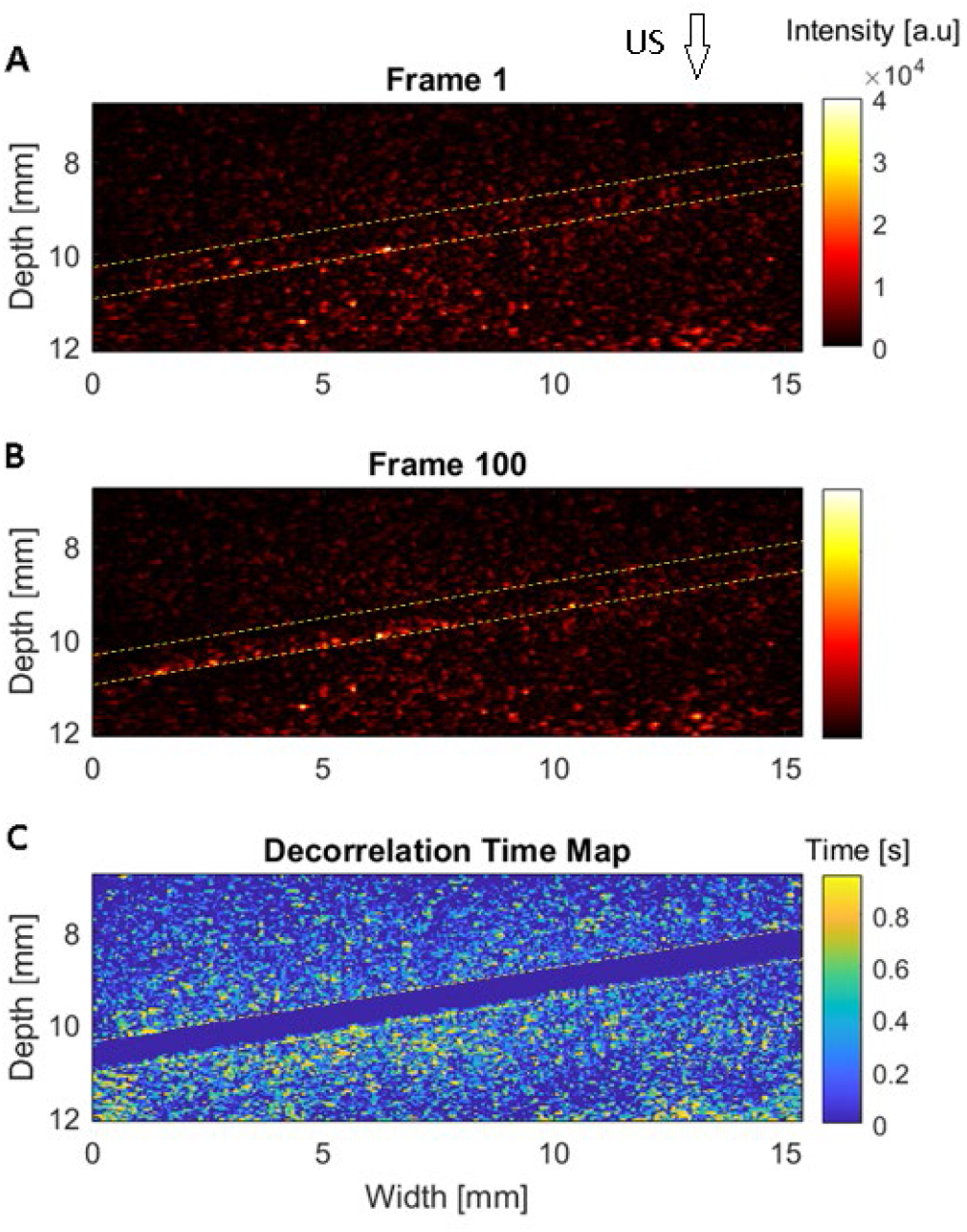
A) First frame of a nonlinear contrast (NLC) video of an agarose gel phantom with nanobubbles (NBs) entrapped in the agarose matrix (∼4×10^9^ NBs) and a matching NB:PBS dilution perfused through a central diagonal channel flowing at 250 µl/min. The filled channel is indistinguishable from surroundings. B) Final frame of the agarose gel phantom in NLC mode. NLC has slightly decreased in surroundings due to bubble destruction. C) Decorrelation time map of the flow phantom. Dashed white lines indicate upper and lower channel boundaries. Arrow indicates direction of the ultrasound beam.

A representative decorrelation map of the flow phantom is shown in Fig. 5-C. DT is higher in the surroundings (up to 1s) where the known entrapped NB population is located. The center channel with flowing agents, however, shows low DT (DT = 0 s). As a result, there is clear delineation between regions of flowing and stationary agents visible in the decorrelation map. Namely, the flow channel is highly distinguishable with low DT while signal from entrapped NBs produce long DT.

### Microfluidic Phantom

To show the sensitivity of decorrelation mapping to agents that are extravasating and diffusing through a collagen matrix, LumenChips were produced with 1.3 μm and 3.7 μm diameter^46^ pore sizes. They were perfused with either NBs or MBs. After initial injection, the devices were continuously imaged for 4000 frames to capture the agent dynamics.

Representative B-mode and NLC mode images are shown at 40, 280 and 680 s time points as a demonstration of the progressive signal increase from agents as they diffused beyond the lumen through the 3.7 μm pore size matrix (Fig. 6-A & C). As shown in Fig. 6, we found that MBs (∼875 ± 280 nm diameter) experienced little diffusion, which is evident by the minimal signal increase beyond the lumen (indicated by the yellow dashed box on NLC images which are outside of the vessel lumen), even after 680 s of continuous flow (Fig. 6-A). Conversely, NB (100-500 nm diameter) B-mode and NLC images show progressive diffusion of NBs beyond the lumen through contrast enhancement with increasing penetration beyond the lumen in time. Like the flow phantom, after penetration of agents into the collagen matrix it was visibly difficult to distinguish between agents in the lumen and those in the matrix from static images.

**Figure 6.**
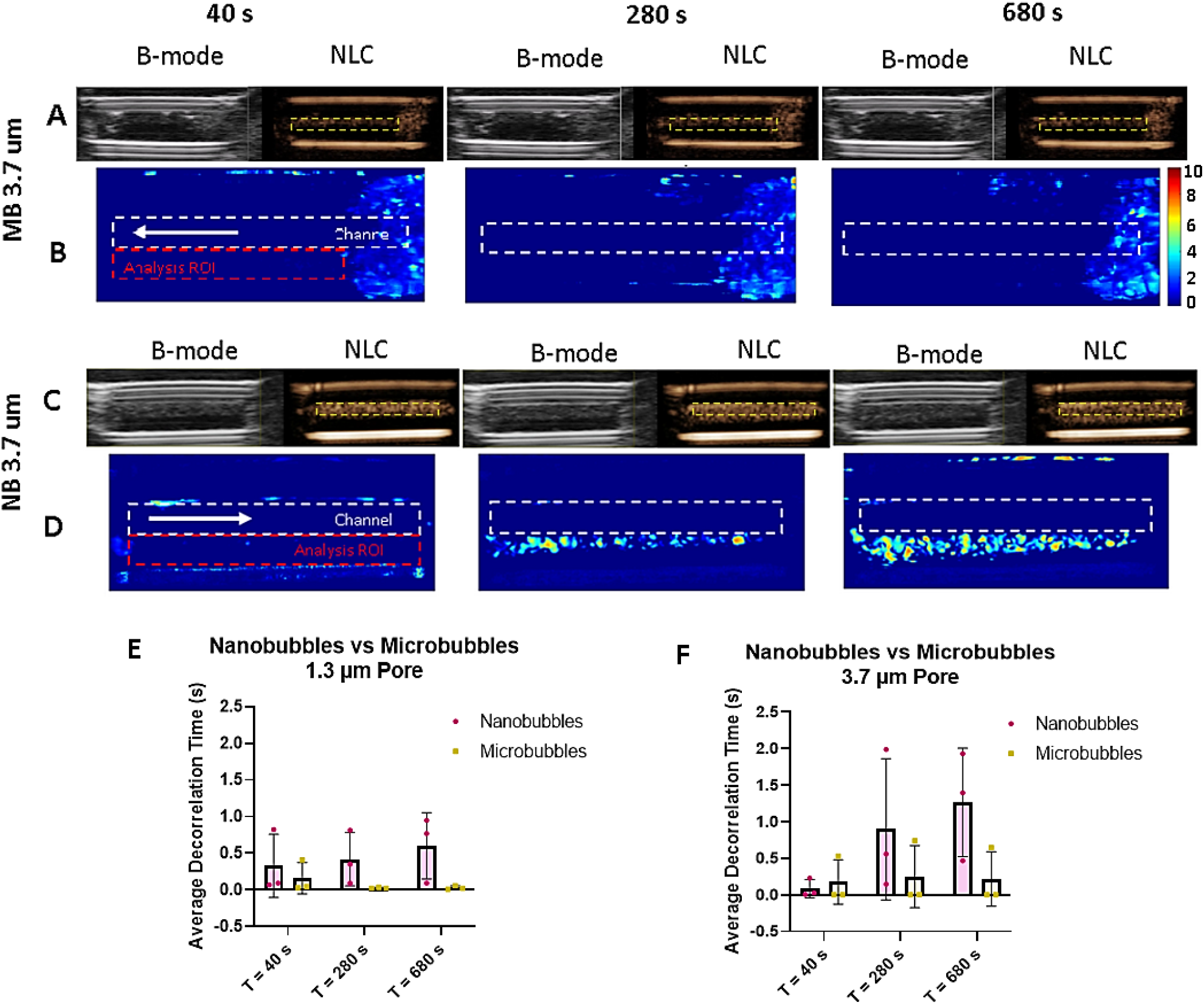
(rows A & C) Representative B-mode and nonlinear contrast (NLC) mode images produced from the VEVO 3100 are shown for microbubbles (MBs) and nanobubbles (NBs) in rows B and D respectively. Decorrelation time maps NLC mode imaging of a microfluidic device containing a collagen I matrix (3.7 µm pore size) chamber with a lumen (0.5 mm diameter) extending through the center of the device. The lumen was injected with either MBs (MB - A & B) or NBs (NB - C & D). Decorrelation time maps were produced from analysis of 200 frames starting from time points of (left to right) 40, 280 and 680 s respectively after NB entrance into the channel. Dashed red ROI indicated area of analysis. White arrow indicates direction of agent flow through the lumen. (E&F) Comparative data for average time-intensity curves from selected regions of interest comparing NBs (fuchsia) to MBs (yellow). Error bars represent standard deviation of averages from 3 trials. E) Decorrelation time averages from collagen matrix with 1.3 µm diameter pore sizes F) Decorrelation time averages from collagen matrix with 3.7 µm average pore sizes.

DT maps were produced from 200 frames at each time point, as shown in Fig. 6-B&D. From decorrelation maps, low DT (0 s) is seen in regions of either no signal or rapidly fluctuating signal such as in the lumen. The DT maps for MBs in the LumenChip did not allow visualization of the lumen – collagen matrix boundary. By comparison, NBs in the 3.7 µm pore matrix produced decorrelation maps with extended DT in the matrix region (up to 10s), corresponding to the increase in signal intensity as NBs diffused through the matrix, which clearly delineate the lumen – collagen matrix boundary. In the 3.7 µm pore matrix, areas with longer DT grew with time when NBs were perfused through the channel (Fig. 6-D) while no increasing area of elevated DT was observed when the lumen was perfused with MBs (Fig. 6-B). Results from the 1.3 µm matrix can be seen in supplemental Fig. S1. With a smaller pore size, MBs and NBs experienced decreased penetration into the extraluminal space.

Results from averaging of the regions of interest (ROIs) from the DT maps depicted in Fig. 6-B&D and Fig. S1 (red dashed rectangles) are shown in Fig. 6 panels E and F. The average DT for MBs, which diffused minimally into the collagen matrix and therefore experienced minimal entrapment, was consistently at 0 s for a pore size of 1.3 µm. NBs that diffused beyond the lumen into the collagen matrix showed elevated average DTs for both collagen concentrations. At 1.3 µm pore size (Fig. 6-E), the DT was consistently greater for NBs than MBs for all imaging windows. In the 3.7 µm pore size (Fig. 6-F) the average DT with NBs was greater than MBs for later times. Overall, in the 3.7 pore matrix average DT of agents increased with a larger pore size while MBs did not result in extended DT.

### Mouse Model

As proof of concept of DT mapping *in vivo*, our analysis was applied to US data from a previous study of a mouse tumour flank model injected with MBs or NBs. Representative images of the first B-mode frame of imaging and then first and final NLC frames of the varying contrast agents are presented in Fig. 7. TICs of the regions of interest are also provided for visualization of the agent wash-in dynamics. Autocorrelation curves of the plotted TICs are also presented in Fig. 7.

**Figure 7.**
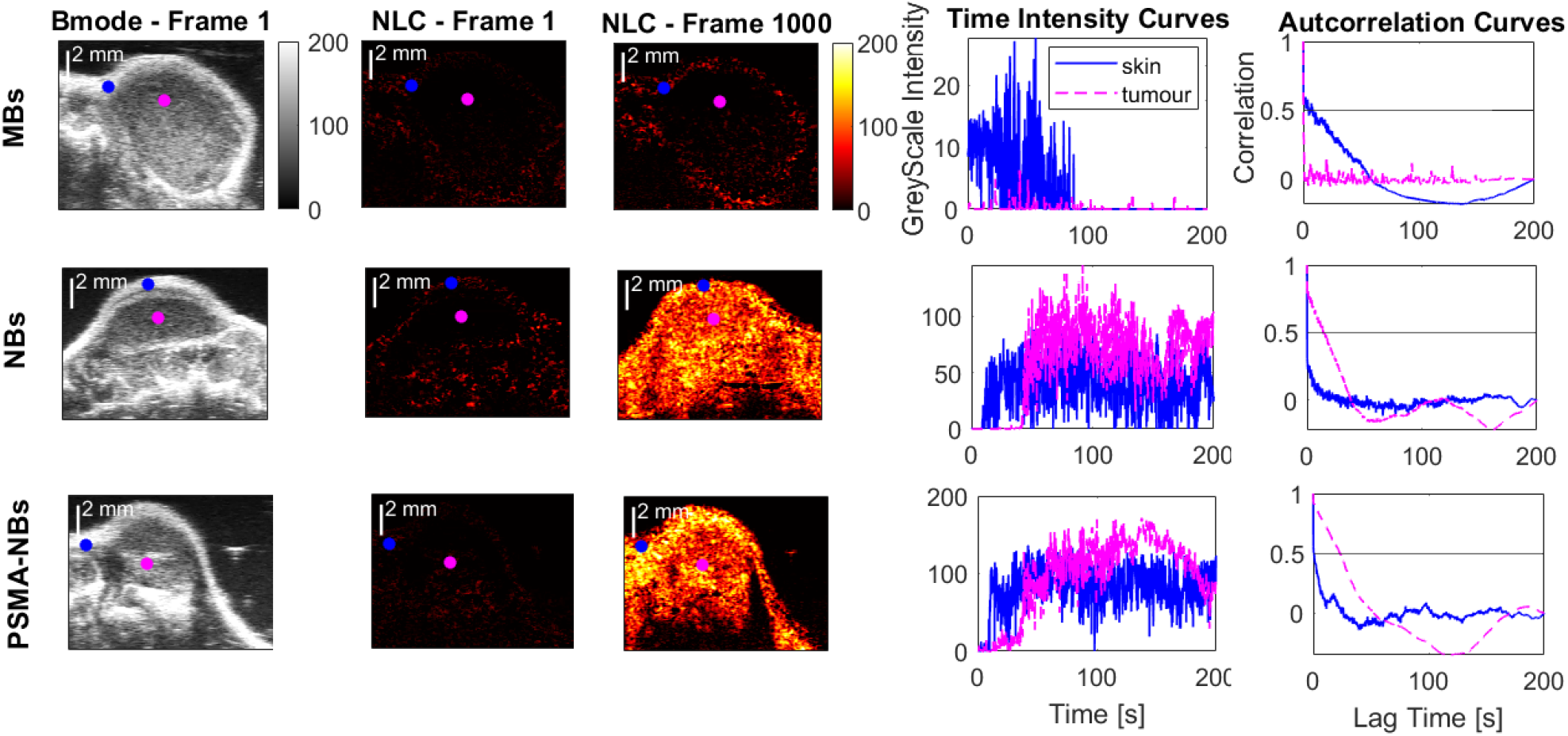
Flank tumour mouse models bearing subcutaneous human PC3-pip tumour xenografts injected via tail vein with either (top to bottom) microbubbles (MBs; Lumason), nanobubbles (NBs), or prostate specific membrane antigen targeted NBs (PSMA-NBs). Representative images (left to right) of: 1) the first frame acquired by B-mode imaging; 2) the first frame acquired by nonlinear contrast (NLC) mode imaging after background subtraction; 3) the final NLC frame after background subtraction; 4) sample time-intensity curves (TICs) from a selected skin (blue) or tumour (fuchsia) locations, as marked in ultrasound images (1-3); and 5) sample autocorrelation curves from skin and tumour selections, as marked in ultrasound images and TICs.

B-mode images of the tumours are shown on the left of Fig. 7 for reference. From the B-mode images, surrounding healthy tissue and skin generate high intensity signal. Relative to surrounding healthy tissue, the tumour cores in B-mode images appear homogeneous with few distinguishable details between any of the models.

NLC data were the focus of this work. As shown in Fig. 7, tumours injected with MBs experienced minimal filling relative to tumours injected with NBs in the first and final frames of the NLC videos. By comparison, tumours injected with the two types of NBs experienced similar tumour filling dynamics.

Consistent with the NLC images shown in Fig. 7, the representative TICs show differences in tumour filling dynamics between MBs and NBs. A transient greyscale intensity increase in NLC mode in the skin regions (dashed blue line) was seen from all agents (NLC Frame 1000 – Fig. 7). In the tumour (pink dashed line) the MBs generated noise-level signal while NBs and PSMA-NBs generated high amplitude signal comparable to the skin with delayed onset and time-dependent signal decrease.

The autocorrelation curves were generated from the pixel TICs. For MBs, where the autocorrelation curve of the tumour dropped from 1 to 0 immediately (Fig. 7. Autocorrelation Curve: MBs - pink dashed curve), the representative autocorrelation curve of the skin TIC (Fig. 7. Autocorrelation Curve: MBs - blue dashed curve) had a more gradual decorrelation. Comparing NBs and PSMA-NBs, autocorrelation curves within the tumour regions showed elevated correlation above 0.5 (50% correlation threshold) for NBs (correlated for 12 s) and the PSMA-NBs (correlated for 20 s) for longer lag time, indicating the source for higher DT.

DT mapping was used to identify prolonged correlation on a pixel-by-pixel basis. DT maps for each tumour injected with either MBs or NBs are presented in Fig. 8-A. Regions of interest from the maps containing or excluding the tumour (See Fig S. 2 for ROI selection) were averaged for broader comparisons between maps. The DT maps and average DTs show restricted access of MBs to deep tumour tissue. Only outlines of the MB tumours were visible (∼1.5 s average decorrelation time in the skin Fig. 8-B) such that areas of prolonged DT (Fig. 8-A) corresponded with NLC signal increase (Fig. 7) indicating the presence of MBs in those regions. By comparison, when NBs or PSMA-NBs were used, the DT was significantly elevated in the tumour core (up to 30 s) showing detailed tumour-core morphology and clear delineation from surroundings. The average DTs for the NB tumours and PSMA-NB tumours ranged from 6-14 s while the average DTs in the skin for the same animals were broadly comparable to those with MBs (1-2 s) as shown in Fig. 8-B. Tumours injected with NBs also showed separation of vessel signal (Fig. 8. D - indicated by red arrows) where high flow rates are to be expected (vessel DT ∼ 0 s vs surrounding DT ∼15 s for NBs and ∼30 s for PSMA-NBs). In the surrounding healthy tissue, the decorrelation time is low, suggesting the lack of NB accumulation.

**Figure 8.**
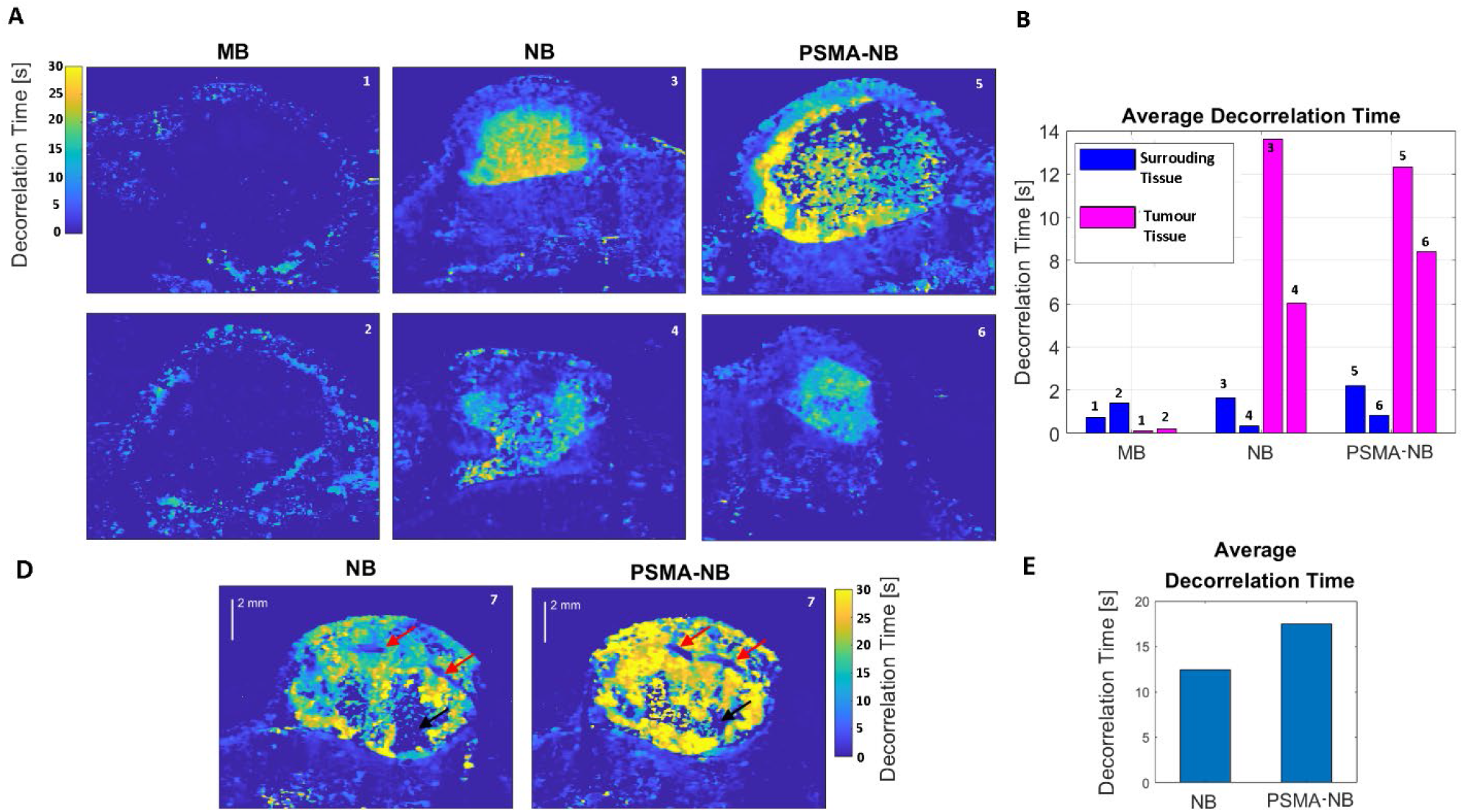
A) Decorrelation time maps of 6 different flank tumour mouse models bearing subcutaneous human PC3pip tumour xenografts that were injected via tail vein with either microbubbles (MBs; Lumason - (left-most column)), nanobubbles (NBs - (center column)), or prostate specific membrane antigen targeted NBs (PSMA-NB; (right-most column)). B) Average decorrelation times from selected skin (blue) and tumour (fuchsia) regions of interest (ROIs) from DT maps. ROIs for averaging shown in supplemental. D) Decorrelation time maps of a single flank tumour mouse model bearing a subcutaneous human PC3pip tumour xenograft that was injected via tail vein with nanobubbles (NBs - left) followed by prostate specific membrane antigen targeted NBs (PSMA - right). E) Average decorrelation times from selected tumour regions of interest for the double injected tumour.

To assess the sensitivity of DT mapping comparing active or passive targeting techniques, DT mapping was applied to a single tumour injected with both NBs and PSMA-NBs at separate times (Fig. 8-C&D). DT increased with active targeting by PSMA-NBs relative to passively targeted NBs in the same tumour. Here, a necrotic region where NBs were not able to access the tumour was identified using DT mapping (Fig. 8-C – NB & PSMA, black arrows). Additionally, a blood vessel running along the imaging plane, as indicated by the red arrows in Fig. 8-D, was clearly delineated from surroundings due to low DT from fast moving NBs. For validation of the vessel location and comparison to NLC images, select NLC frames from the tumour displayed in Fig. 8-C are provided in Fig. S3.

## DISCUSSION

Currently there are no effective image-processing techniques to extract information from dynamic nanobubble-contrast enhanced US scans that identify perfused areas and provide maps of localized agent accumulation. Imaging the EPR effect *in vivo* has been achieved through tumour selective imaging using fluorophore-conjugated nanoprobes and an optical *in vivo* imaging system (IVIS^®^).^2,49^ However, to image the probe accumulation, tumour-bearing mice were sacrificed and their tumours were removed.^2^ Gold nanoparticles (GNPs) have been proposed as a contrast agent for photothermal molecular imaging of cancer^50^, but optical-based methods typically suffer from low penetration depth and are therefore limited in their applications. Ultimately, real-time imaging techniques for the diagnosis and characterization of EPR are sought. These techniques could enable clinicians to preselect patients with high EPR-tumours who are more likely to respond to nanotherapeutic treatments.^6,13^ For such an imaging technique to be successful and lead to clinical translation, it must meet several criteria: (1) high sensitivity to key identifiers of EPR (vascular permeability, vascular structure, lymphatic drainage etc.), (2) relatively low-cost, (3) strong safety profile, and (4) rapid, near real-time output. Clinical translation would be accelerated by building on existing technology and protocols. The most rigorous techniques for identifying adhesion and accumulation of targeted agents require invasive or destructive protocols, requiring excision. Many of the non-invasive techniques that are available use ionizing radiation, which contributes to a low safety profile. The remaining techniques which are non-invasive and do not use ionizing radiation are expensive technologies, requiring large investments in money, time, and skilled labour and are therefore limited in accessibility and impact. The aim of the DT mapping technique was to develop a safe, minimally invasive, and inexpensive technology producing detailed images that utilize NB-specific tumour biomarkers. The power of such a technique lies in the broad applicability across vascular diseases where elevated vascular permeability is a common pathology. Contrast enhanced US imaging has long met the requirements of a safe, non-invasive imaging technology that is accessible. Here, the decorrelation of the NLC echo signals from NBs in tumours converts large datasets into single images, which capture the details of the agent dynamics inside and outside of the tumours.

For the signal of a pixel in NLC to correlate in time, the signal sampling rate needs to be greater than the timescale of changes in the NLC TICs. NLC images result from the isolated detection of nonlinear scatter from contrast agents. Thus, changes in the NLC intensity within an imaging voxel could be quantified using signal correlation if the timescales related to the changes are lower than the US imaging framerate. Conversely, NLC intensity fluctuations that are more rapid than the frame rate are not detected due to temporal resolution limits and therefore do not contribute to the signal correlation. NLC intensity fluctuations are caused by agent motion through the voxel (e.g. by flow), agent destruction (e.g. by dissolution or burst), or tissue motion (e.g. from heart beat or breathing). Thus, for a fixed voxel size, the sampling rate (framerate) determines the sensitivity threshold to different agent dynamics.

The highest framerate we explored was used to image our flow phantom setup (20 fps). At this frame rate, a scattering source would have to remain in the imaging voxel for at least 0.05 seconds to produce a non-zero DT. Using the pixel dimensions (width = 0.045 mm, height = 0.02 mm) as an approximation of the voxel dimensions, for a stable agent to be in the voxel for at least 0.05 s, it would be moving at a velocity of ∼1 mm/s or slower. This can be thought of as a velocity threshold for non-zero DT. However, in the flow phantom setup, NB agents in the flow channel were moving at a velocity of approximately 20.4 mm/s (0.51 mm channel diameter; volumetric flow rate = 250 µL/min; velocity calculated from volumetric flow rate), a factor of 20 over the velocity limit for non-zero DT. Thus, agents in the channel space were moving too fast, leading to 0-s DT (Fig. 5-C). For reference, red blood cell velocities have been measured ranging from as low as 0.5 mm/s (cerebral capillaries) to as high as 52–152 cm/s (aorta). ^52,53^ For the same pixel dimensions as mentioned above, a framerate of 5 fps would lead to a non-zero DT velocity threshold of 0.25 mm/s. Even the lowest red blood cell velocities would be too fast to correlate signal from one frame to the next leading to extremely low DTs (0 s).

By comparison, agents embedded in the phantom matrix were trapped in a single voxel space for longer than the inter-frame period and therefore corresponded to longer decorrelation times. Autocorrelation analysis has previously been applied to assess the motion of agents in agarose using US^40^. With sufficiently high agarose concentrations, NB agents with diameter equal to or greater than the diameter of the agarose pores become trapped and produce flattened autocorrelation curves (increased decorrelation time), indicating restricted motion^40^. In the current work, the concentration of agarose from the previous study, which corresponded with hindered NB motion (1.25% w/v agarose in PBS^40^), was used to establish a population of entrapped agents. Consistent with the previous findings, ^40^ the regions of our nanobubbles entrapped in the agarose phantom corresponded with locations of prolonged DT (up to 1s - Fig. 5-C). Thus, the imaging framerate controls the sensitivity of the mapping technique to motion and allows the separation between blood pool and extravascular agents. Consistent with applications of US echo decorrelation analysis to flow velocimetry, agents in the flow channel yielded significantly lower DT compared to regions with entrapped agents. Unlike other studies, here the use of low framerate for data acquisition was used as an inherent flow-velocity threshold, to limit observations to stationary and near-stationary signals.

In the presence of increased vascular permeability and extravasation, bubble entrapment in the extraluminal space increases over time. We used a microfluidic platform, LumenChip, with pore sizes of 1.3 and 3.7 µm on average^46^ to mimic the permeability and extravasation kinetics of the leaky tumour vasculature. Endothelial gap diameters have been shown ranging from 0.2 to 1.4 µm in venules after action of substance P or bradykinin^51^, which is important in chronic inflammatory response. Intercellular gaps in tumour vascular systems usually range from 100–800 nm^52^ and can be as large as 1.2 µm in diameter. The pore sizes explored here are of the same size of some pathologically large endothelial gaps, especially for cancers like mammary carcinoma,^53^ and are therefore clinically relevant. Our results further exemplify the physical limitations of larger bubble agents for pathologies with endothelial gaps smaller than the pore size explored here. Diffusion by a nano-agent in a polymer solution is related to the viscosity of the polymer solution, the size of the particle, and the flexibility of the probe particle.^54^ In our microfluidic model, diffusion of the MBs or NBs, when it occurred, was axial to the US propagation path as a result of acoustic radiation force, which has been established as a method to increase access of agents to tissue beyond the blood pool. The role of acoustic radiation force on the agents on this microfluidic setup has been investigated in depth previously.^55^

As with the flow phantom, in the LumenChip (collagen matrix in a microfluidic device), we showed that after extravasation, the velocity of agents can be slowed sufficiently for sensitive detection by DT mapping. The luminal agents were moving at a velocity (∼1.06 mm/s; flowrate = 50 µL/min) above the non-zero DT threshold (0.08 mm/s; 5 fps; pixel height = 0.0176 mm), leading to 0 s DT in the channel (Fig. 6). Meanwhile, bubbles of small enough diameter were able to travel beyond the luminal space, as is depicted in B-mode and NLC images from Fig. 6 comparing MBs to NBs. Signal outside of the lumen corresponded with prolonged DT (up to 10 s) due to the slowing of agents caused by collisions with the matrix. We observed prolonged DT in a larger area as time and diffusion progressed (Fig. 6).

Interestingly, a discrepancy was seen between the maximum DTs of agents in the microfluidic device compared to the flow phantom. Agents in the microfluidic device reaching the collagen matrix reached a maximum DT ten times greater (10 s) than those immobilized in the flow phantom (1 s). The reason for the extended DT is not clear but may be related to stability of contrast agents in the acoustic field. As agents are subject to prolonged US exposure, they are destabilized and dissipate due to gas diffusion.^56^ Elevated shear stresses (e.g., by turbulent flow), increased ambient pressure, increased temperature, and low pH are all factors that can destabilize an agent population and can occur *in vivo*. ^57^ Shielding agents from fluctuations in any of these factors can have stabilizing effects. Additionally, uptake of agents into cells (e.g., via endocytosis) can have a stabilizing effect due to shielding of the agent from incident US stimulation.^41^ Without bubble destruction, decorrelation time would be elevated for an agent due to prolonged scatter potential.

Finally, DT mapping for cancer detection using targeted and untargeted NBs was then tested *in vivo* to demonstrate proof of concept in a clinically relevant scenario. Due to the ease of clinical translation offered by commercially available and FDA approved agents, MBs have also been a major focus in US molecular imaging.^58,59 19–22^ However, their direct access to cancer cells is limited due to their size. ^60^ Contrast enhancement in the tumour by NBs was higher than MBs, showing demonstrable tumour filling and enhanced deep-tissue access (Figs. 7&8). Average DT was consistent across all models when analyzing skin and non-tumour regions (∼1 s). MB and NB enhancement in the highly vascularized tumour periphery was also similar. When evaluating the tumour core, the differences in contrast dynamics were apparent between MBs and NBs. DT from MB-enhanced scans was near zero due to the lack of signal from MBs. Both targeted and untargeted NBs produced average tumour decorrelation times up to 7 times higher than the surrounding tissue (Fig. 8). NBs showed delayed access relative to skin, an indicator of longer transit time (Fig. 7 - TICs NBs and PSMA-NBs). NB TIC curves also exhibited a decrease in signal intensity over time, which is an indicator of agent destruction from prolonged US exposure without readily available agents to replace them. Overall, the high level of detail and contrast offered by the DT maps shown in Fig. 8 is likely due to the prolonged presence of NBs as they extravasate from the vasculature and slow in the tumour core.

In a single snapshot comparing untargeted to targeted agents, the DT map showed a complete outline of the tumour. The high rate of speckle fluctuation in a vessel region (visible from NLC– see Fig. S3) is readily visible in the DT map (Fig. 8-C red arrow). The diameter of the vessel was determined to be approximately 350 µm. A vessel of this size would be expected to have a flow rate similar to a large capillary (0.3 mm/s) ^60^. At a frame rate of 5 fps with a pixel height of 0.03 mm, the fastest moving NB in the field of view resulting in non-zero decorrelation time had to be moving at 0.067 mm/s, an order of magnitude slower than the expected flow rate of the indicated vessel. Therefore, the agents in the tumour region were significantly slower than the blood pool. DT is sensitive to patterned motion (e.g., periodic motion from breathing or pulse), which contributes to the visualization of the heterogeneity of the mouse tumour as seen in Fig. 8-C.

The study has some limitations. Although image registration was applied to minimize motion effects on pixel signals, ill-matched frames and residual artifacts were likely still present and result in error in measured DTs. Locations of zero signal enhancement resulted in a DT of 0 s. Other groups have normalized their decorrelation values to aid in the discrepancy between pixels, which had no signal and those which had signal which decorrelated rapidly^33^. Future applications of this technique should implement such a normalization to help with the identification of blood vessels perpendicular to the field of view and the identification of necrotic tissue.

## CONCLUSIONS

This work presents the first use of DT mapping as a NB-specific tool for extraction of dynamic contrast enhanced US information. DT calculated from the autocorrelation of TICs from NB contrast enhanced US results from: (1) agent velocity, (2) agent stability, and (3) *in vivo* motion. The methods presented here rely on the small size and high echogenicity of NBs in NLC, leading to extravasation and contrast enhancement dynamics not achievable with MBs. We showed the successful application of ultrasound echo decorrelation for *in vitro* imaging of NB US contrast agents and *in vivo* prostate cancer models using the same US contrast agents. High contrast images were produced that rapidly show the locations of prolonged agent signal, consistent with localized NB accumulation. Despite the potential offered by EPR for nanoagents, the heterogeneity of EPR remains to be one of the greatest hurdles for EPR-based nanotherapeutic strategies. Overall, the EPR effect can differ between individuals, cancers, tumours,^61^ and even between tumour regions.^62^ The application of DT mapping to clinical diagnostics would enable rapid assessment of stagnated NB movement when enhanced vascular permeability and retention (EPR) is present. Because this new biomarker is sensitive to agent dynamics, applications include but are not limited to cancer diagnosis. Elevated levels of vascular permeability also occur during inflammation and is a hallmark of many inflammatory diseases (e.g. diabetes).^63^ Improvements for cancer diagnosis include improved early detection, increased specificity of US molecular imaging when using molecular targeting and enhanced tumour margin delineation. The sensitivity of the DT method on a pixel-wise basis, when applied to the NB contrast enhanced US data, allows for spatial mapping of the heterogeneity of vascular permeability. Application of DT mapping of NB signal to tumours could lead to a simple tool for examining tumour susceptibility to nanotherapeutics,^12,13^, and could mean rapid diagnostics and down stream patient preselection^6^ and personalized cancer therapy.

## Acknowledgments & Funding

This work was funded by and supported in part by the National Institutes of Health (R01EB025741, T32GM007250, F30HL160111). This research was also supported by the National Institute of General Medical Sciences (T32GM007250); the National Center for Advancing Translational Sciences (TL1 TR000441); the National Heart, Lung, and Blood Institute (T32HL134622, F30HL160111, R01HL133574, and R42HL160384); the National Institute of Biomedical Imaging and Bioengineering (R01EB025741 and R01EB028144); the National Cancer Institute (R21CA253108); and National Science Foundation (NSF) CAREER Award 1552782. Figures created with BioRender.com. The authors would like to thank F. M. Berg for review of this manuscript.

## Notes

### Competing Interest Statement

The authors have declared no competing interest.

