## Supplemental Materials for "Decorrelation Time Mapping as an Analysis Tool for Nanobubble-Based Contrast Enhanced Ultrasound Imaging"

### Supplementary Material

The following data are provided to offer the reader supporting information to the methods presented.

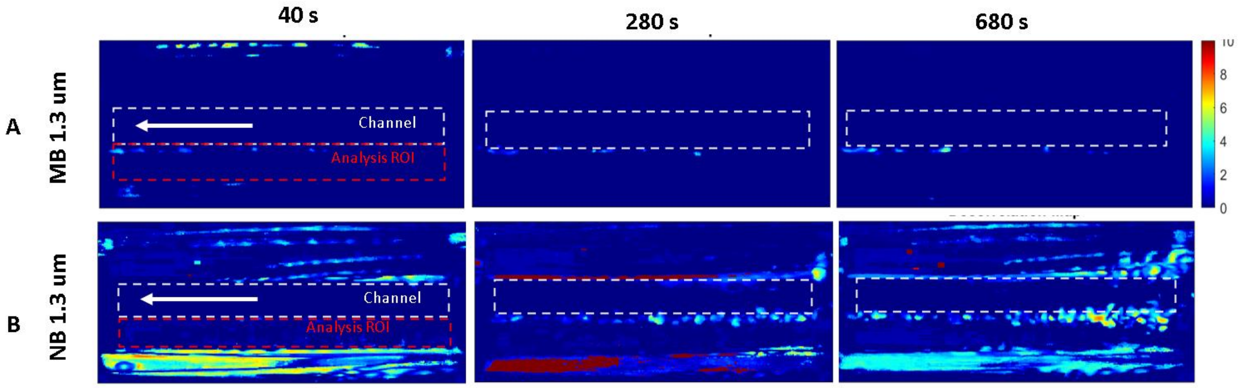


**Fig. S1.** Decorrelation time maps from nonlinear contrast (NLC) mode imaging of a microfluidic device containing a collagen I matrix (1.3 µm pore size) chamber with a lumen (0.5 mm diameter) extending through the center of the device. The lumen was injected with either microbubbles (MB – A) or nanobubbles (NB - B). Decorrelation time maps were produced from analysis of 200 frames starting from time points of (left to right) 40, 280 and 680 s respectively after NB entrance into the channel. Dashed red ROI indicates area of analysis. White arrow indicates direction of agent perfusion through the lumen.

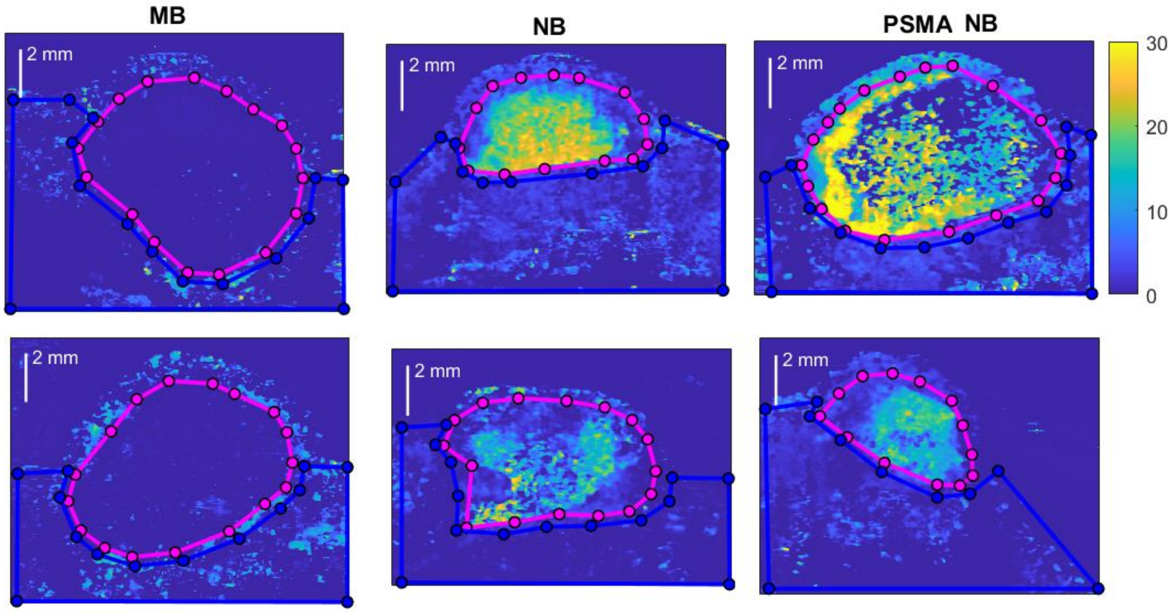


**Fig. S2.** Decorrelation time maps of flank tumour mouse models bearing subcutaneous human PC3pip tumor xenografts that were injected via tail vein with either microbubbles (MBs; Lumason - (left-most column)), nanobubbles (NBs - (center column)), or prostate specific membrane antigen targeted NBs (PSMA-NB; (right-most column)). Average decorrelation times were determined from skin ROIs (blue outline) and tumour ROIs (fuchsia).

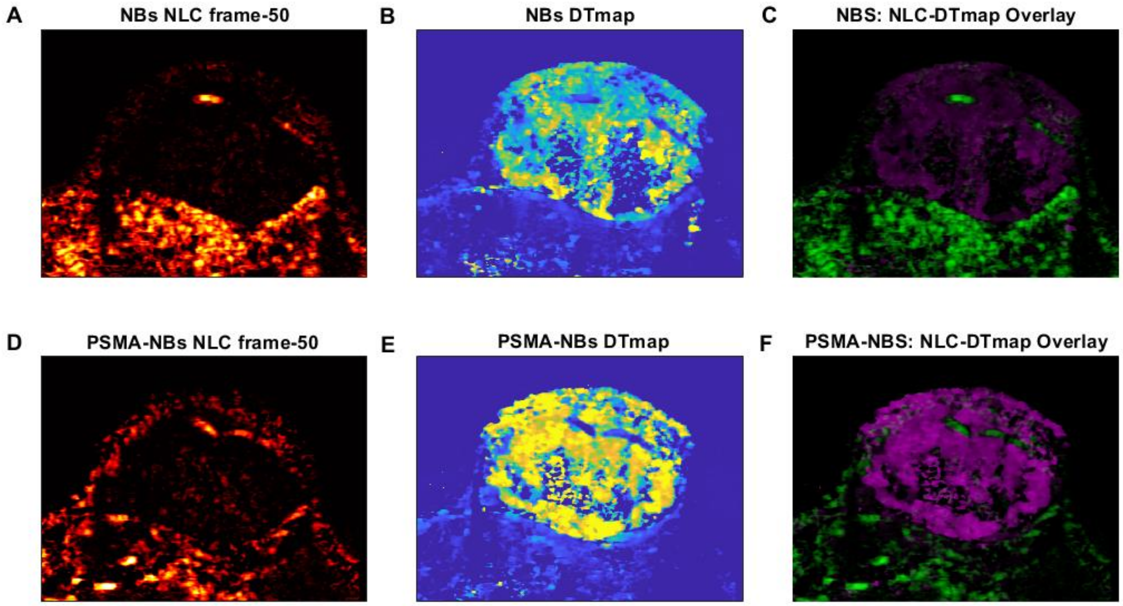


**Fig. S3.**A single flank tumour mouse model bearing a subcutaneous human PC3pip tumor xenograft that was injected via tail vein with prostate specific membrane antigen nanobubbles (PSMA-NBs) followed by plain NBS. A) 50$^{th}$ frame of NLC image from tumour injected with NBs which highlights a large vessel through the tumour region. B) Decorrelation time map of tumour injected with NBs. C) Overlay of NLC image with DTmap shows sensitivity of DT mapping to flow regions and the suppression of signal from established blood pool agents. D) 50$^{th}$ frame of NLC image from tumour injected with PSMA-NBs which highlights two blood vessels through the tumour region. B) Decorrelation time map of tumour injected with PSMA-NBs. C) Overlay of NLC image with DTmap shows sensitivity of DT mapping to flow regions and the suppression of signal from established blood pool agents in the presence of active targeting.
